# Mg^2+^ Sensing by an RNA Fragment: Role of Mg^2+^ Coordinated Water Molecules

**DOI:** 10.1101/2020.06.04.133371

**Authors:** Antarip Halder, Sunil Kumar, Omar Valsson, Govardhan Reddy

## Abstract

RNA molecules selectively bind to specific metal ions to populate their functional active states making it important to understand their source of ion selectivity. In large RNA systems, metal ions interact with the RNA at multiple locations making it difficult to decipher the precise role of ions in folding. To overcome this complexity, we studied the role of different metal ions (Mg^2+^, Ca^2+^ and K^+^) in the folding of a small RNA hairpin motif (5′-ucCAAAga-3′) using unbiased all-atom molecular dynamics simulations. The advantage in studying this small system is that it requires specific binding of a single metal ion to fold to its native state. We find that even for this small RNA, the folding free energy surface (FES) is multidimensional as different metal ions present in the solution can simultaneously facilitate folding. The FES shows that specific binding of a metal ion is indispensable for its folding. We further show that in addition to the negatively charged phosphate groups, spatial organization of electronegative nucleobase atoms drive the site specific binding of the metal ion. Even though the binding site cannot discriminate between different metal ions, RNA folds efficiently only in Mg^2+^ solution. We show that the rigid network of Mg^2+^ coordinated water molecules facilitate the formation of important interactions in the transition state. The other metal ions such as K^+^ and Ca^2+^ cannot facilitate the formation of such interactions. These results allow us to hypothesize possible metal sensing mechanisms in large metallo-riboswitches and they also provide useful insights for the design of appropriate collective variables for studying large RNA molecules using enhanced sampling methods.

## Introduction

Ribonucleic acid (RNA) molecules fold into competent structures to participate in a plethora of cellular processes such as protein synthesis,^1^ catalysis,^2^ regulation of gene expression,^3,4^ gene editing,^5,6^ etc. Metal ions are inextricably linked to the assembly of functional RNA structures.^7–9^ Metal ions renormalize the negative charge on the phosphate groups and decrease the persistence length of the RNA chain enabling the chain to collapse in the initial stages of folding.^10–12^ Metal ions are also found to affect the rate of conformational rearrangements, ^13–15^ modify the kinetic free energy barriers, ^16^ participate in the formation of transition states,^17^ alter the dominant folding pathways^18–20^ and are shown to be intricately involved in the catalytic mechanism of RNAs such as ribozymes.^21–23^

Mg^2+^ is the most ubiquitous metal ion, which is bound to the RNA in crystal structures.^24^ Mg^2+^ in RNA folding are classified into two types: diffuse and site-binding, depending on its contribution to the stabilization of the RNA structure.^25^ The diffusely bound Mg^2+^ binds to the polyanionic RNA chain nonspecifically and result in the overall collapse of the chain dimensions. Whereas, the site-bound Mg^2+^ is attracted to specially arranged binding pockets and aid in the stabilization of secondary and tertiary structures. Although recent simulations^26^ on *Azoarcus* ribozyme show that Mg^2+^ condensation on the RNA chain is site specific and depends on the structure of the folded state.

There is an extensive effort to understand how metal ions influence RNA folding using various biophysical methods.^27–31^ Due to the fluctuating nature of the ion atmosphere around the RNA, it is extremely difficult to study its influence on RNA folding using traditional techniques, which probe structure such as X-ray crystallography, NMR, or (cryo-) electron microscopy. These techniques only identify ions, which specifically bind to a group of RNA atoms as a part of their coordination shell and stabilize the RNA structure.^32^ Theoretical and computational techniques are playing an important role in complementing experiments to identify the role of ions in various other aspects of RNA folding.^33^

Molecular dynamics (MD) simulations using atomistic^34–38^ and coarse-grained^39–46^ force fields have provided important insights into RNA folding. Nevertheless, a comprehensive understanding of how large RNAs selectively recognize specific metal ions and fold to their respective active conformations is far from complete. For example, metal ion sensing riboswitches^47,48^ recognize a specific type of metal ion (Mg^2+^,^49^ Mn^2+^,^50,51^ Co^2+^,^52^ Ni^2+^,^52^ etc.) and depending on the metal ion concentration, they switch from unfolded to folded conformations to regulate the downstream gene expression. However, the source of selectivity towards specific ions in this type of large RNA systems (*>* 90 nt) is not understood at the molecular level. In large RNA systems, metal ions interact with the RNA at multiple locations making it difficult to decipher the precise role of site-bound ions in folding. To overcome this complexity and probe the source of metal ion specificity in RNA, we have identified a simple yet robust RNA tetraloop system that requires a single site bound metal ion for its folding and studied its unfolding/refolding mechanism using atomistic MD simulations.

RNA tetraloops are small three-dimensional hairpin motifs that are found ubiquitously in almost all functional RNAs and are associated with important structural and functional roles.^53^ A tetraloop hairpin motif is composed of an A-form helical stem capped by four nucleotides arranged in a specific three-dimensional structure (Figure 1). Due to their small sizes and availability of ample experimental data, the tetraloops are considered to be ideal testbed for atomistic MD simulation studies on RNA.^33^ There are only a limited number of nucleotide sequences that form these tetraloops and a majority of the known tetraloops either belong to GNRA or UNCG tetraloop category, where N is any residue and R is any purine residue (A/G).^54^ Folded structures of RNA hairpin motifs with GNRA (or UNCG) tetraloop sequences are conserved and are thermally highly stable as the bases in the loops display ordered stacking interactions in addition to specific non-Watson-Crick base-pairs and backbone conformations.^55^

**Figure 1:**
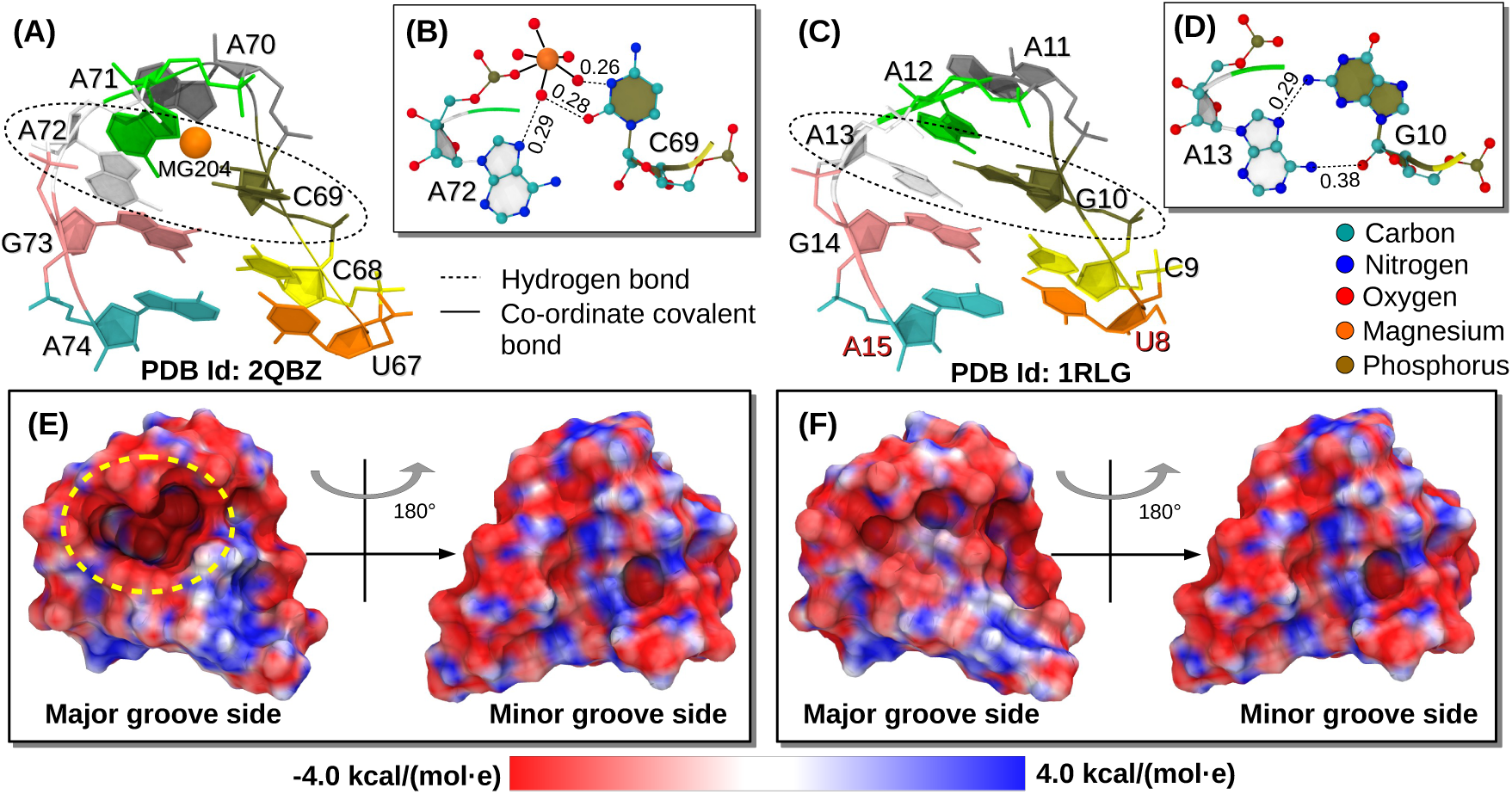
(A) Crystal structure^49^ of the hairpin motif containing an A-form helix capped by the L4 tetraloop. The Mg^2+^ present inside the loop is shown in orange color. (B) The Mg^2+^ has water mediated interaction with C69 and A72 nucleobases. Hydrogen bonds are shown as broken lines in black. The interatomic distances between the hydrogen bond donor and acceptor atoms are given in nm. (C) Crystal structure of a GNRA tetraloop hairpin motif containing the same A-form helix. The terminal residues of the motif are manually edited (C8 → U8 & G15 → A15) to resemble the former hairpin motif. (D) Closing residues of the tetraloop form a G10:A13 *trans*S:H pair. Electrostatic potential surfaces of the hairpin motif containing the (E) L4 tetraloop and (F) GNRA tetraloop, respectively with views from the major and minor groove side. An anionic cavity present at the center of the L4 tetraloop is visible from the major groove side (encircled with a yellow broken line). No such cation binding cavity is present in the GNRA tetraloop. The electrostatic potential surfaces are calculated by solving the Poisson-Boltzmann equation^56^ using the PBEQ Solver server^57^ present in CHARMM-GUI.^58^ The parameters used for the calculation are given in the SI.

In this work, we studied the folding of a special RNA tetraloop (L4 loop in M-box riboswitch^49^), which despite not having the required sequence, folds into the shape of a GNRA tetraloop (Figure 1A). The L4 tetraloop is part of the magnesium sensing M-box riboswitch that switches conformations with the change in Mg^2+^ concentration and regulates downstream gene expression via termination of the transcription process. Crystal structure^49^ of magnesium bound folded aptamer domain (PDB ID: 2QBZ) shows that a Mg^2+^ (Mg204) is present inside the loop. This Mg^2+^ holds together the loop-closing bases (C69 and A72) via its coordinated water molecules (Figure 1B), and facilitates the L4 loop to fold into the shape of a conventional GNRA tetraloop (Figure 1C). This Mg^2+^ mediated inter-nucleotide interaction compensates for the signature G:A *trans*S:H base pair of a conventional GNRA tetraloop (Figure 1D). Electrostatic potential surface for the folded conformation of a L4 tetraloop hairpin motif shows the presence of an anionic cavity at its center (Figure 1E). This is absent in the conventional GNRA tetraloop hairpin motif (Figure 1F). The anionic cavity facilitates cation binding enhancing the stability of the folded conformation, but the source of specificity towards Mg^2+^ remains ambiguous. The simple L4 tetraloop system with a single site for metal cation binding is an ideal system to probe the source of specificity towards selective metal ions in RNA fragments and also study the mechanism of tetraloop folding.

We have carried out all-atom MD simulations of L4 tetraloop hairpin motif in different metal ion concentrations and show that the binding of a metal ion is indispensable for its folding. The site specific binding of the metal ion is guided by the spatial organization of electronegative nucleobase atoms. However, the binding site cannot discriminate between different metal ions. We find that in a mixture of monovalent and divalent ions, the corresponding free energy surface (FES) becomes multi-dimensional as both the ions contribute towards folding. We designed appropriate collective variables (CVs) to compute the FES, which describe the complex folding mechanism of the L4 tetraloop. The FESs show that the L4 tetraloop folds efficiently only in Mg^2+^ solution. Further, the structures in the transition state ensemble (TSE) reveal that a hydrated Mg^2+^ is involved in the transition state (TS) where it plays a critical role in stabilizing the loop part of the motif. The subsequent formation of the A-form helix also requires crucial contribution from the Mg^2+^ coordinated water molecules. We argue that in the absence of rigid octahedral geometry of the water molecules in the first coordination shell of the metal ion, RNA cannot efficiently fold to the native state.

## Materials and Methods

Enhanced sampling methods have been used to study the folding of GNRA and UNGC loops along with other RNA hairpin motifs.^34,35,59,60^ These simulations showed that due to the drawbacks in the forcefields, the native conformations of tetraloops obtained from the crystal structures and NMR, do not correspond to the global minimum in the FES. MD simulations using replica-exchange technique reported folding of UNCG and GNRA tetraloops to their respective native structures.^59,61,62^ However, they are only accounted for a small fraction of the conformations populated (∼ 10% in Chen-García and < 1% in AMBER ff14 forcefields, respectively) in the ensemble with the lowest-temperature, *T* = 277 K.

Recent modifications to the AMBER ff14 force field made by Tan and co-workers alleviated some of the problems in the forcefields to study tetraloop folding.^63^ In the simulations using this modified forcefield,^63^ the population of the native tetraloop conformations increased to 40% in the ensemble with the lowest-temperature *T* = 280 K. Two other recent versions of AMBER force field promise further improvements in sampling the folded state of RNA tetraloops.^64,65^ These improvements in the forcefields will facilitate the study of the role of metal ions in the tetraloop hairpin motif formation using atomistic MD simulations.

### MD simulations

In this work we have used GROMACS simulation program (version 5.1.2)^66,67^ and a modified AMBER ff14 force field^63^ along with TIP4P-D water model^68^ to study the role of different metal ions on RNA tetraloop folding. It is challenging for a force field to capture all the important physical properties of RNA.^33^ The modified AMBER ff14 force field was shown to sample the RNA tetraloop conformations reasonably well^63^ despite its drawbacks.^64^ The folded structure of the eight nucleotide long L4 tetraloop hairpin motif (U67 to A74) is taken from the X-ray crystal structure of M-box riboswitch (PDB ID: 2QBZ^49^). The sequence of the hairpin motif is U^67^CCAAAGA^74^ and the phosphate group at the 5′-end is removed. The hairpin is placed at the center of a cubic box of dimensions ≈ 6.3 nm, and is solvated with TIP4P-D^68^ water molecules. Periodic boundary conditions are applied in all the directions. The negative charges present on the RNA backbone are neutralized by adding seven K^+^ ions. This system corresponds to [K^+^] ≈ 45 mM and [Mg^2+^] = 0 mM. To study the role of Mg^2+^ and Ca^2+^ on the tetraloop folding, we prepared two additional systems, where the concentration of Mg^2+^ and Ca^2+^ is increased to 65 mM, respectively. To these systems appropriate number of Cl^−^ ions are added to maintain charge neutrality. Further details of these three systems are given in Table 1. The simulation box is initially subjected to energy minimization using steepest descent minimization algorithm with an energy cut-off of 1000 kJ/mol/nm.

**Table 1:**
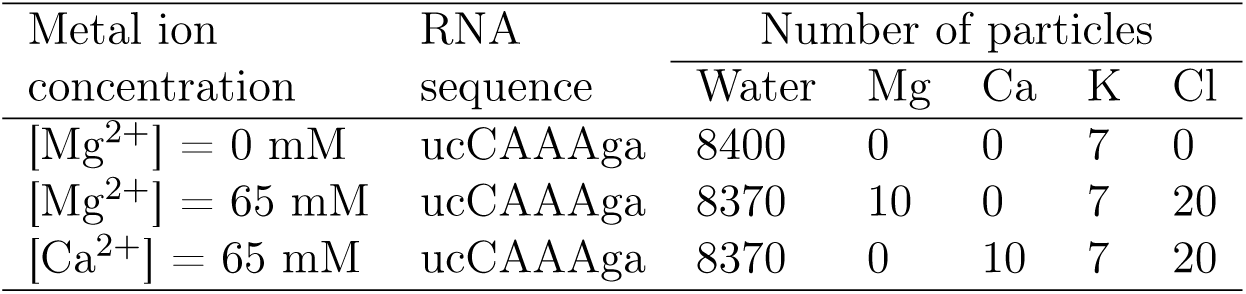
Description of the three L4 tetraloop systems studied

The energy minimized systems are equilibrated in three steps: (i) position restraints are applied on the RNA atoms and ions using a harmonic potential with a force constant of 1000 kJ/(mol nm^2^) and only the solvent molecules are equilibrated for 2 ns in the NVT ensemble, (ii) position restraints are applied on the RNA atoms and ions and only the solvent molecules are equilibrated for 4 ns in the NPT ensemble and (iii) position restraints are applied only on the RNA atoms and both the solvent molecules and ions are equilibrated for 4 ns in the NPT ensemble. Finally, the equilibrated systems are subjected to production runs. The temperature *T* is maintained at 300 K using the stochastic velocity rescaling thermostat developed by Bussi *et al*.^69^ (with time constant *τ*_*t*_ = 0.1 ps) and the pressure *P* is maintained at 1 atm using Parrinello-Rahman barostat^70^ (with time constant *τ*_*p*_ = 2.0 ps). During the production run, all the bond lengths involving the hydrogen atoms are constrained using the LINCS algorithm.^71^ The simulation is advanced using the leap-frog integrator with a 2 fs time step. The long-range electrostatic interactions are computed using particle mesh Ewald summation (PME) method^72^ using 1 nm cutoff for both electrostatic and van der Waals interactions. For each system, we ran two independent trajectories each of length ≈ 10 *µ*s long, and the cumulative length of all the trajectories is ≈ 60 *µ*s (Figure S1). We saved the simulation trajectory every 5 ps for analysis. The first 500 ns of data from each trajectory are ignored and the rest of the data are used for analysis.

### Computing the FES

The FES corresponding to the tetraloop folding is calculated using *G*(**s**) = −*k*_B_*T* ln(*P* (**s**)), where *P* (**s**) is the probability distribution of the set of CVs **s**, and *k*_B_ is the Boltzmann constant. We estimated *P* (**s**) by histogramming the values of the CVs **s** computed from the simulation data. The FESs are plotted using matplotlib tool. ^73^ To probe the contribution of metal ions on folding of the hairpin motif, the free energy is projected onto various CVs viz. eRMSD, end-to-end distance (R_*ee*_), coordination number between metal ions and selected atoms of the RNA fragment (CN_*K*_, CN_*Mg*_ & CN_*Ca*_).

eRMSD is a metric for calculating the deviation between 2 RNA structures.^74^ Unlike the conventional root mean square deviation (RMSD), which use the positions of all nucleic acid atoms to compute the deviation between 2 structures, eRMSD considers only the relative positions and orientations of nucleobases. eRMSD is shown to be more effective and less ambiguous descriptor compared to RMSD. ^34,75,76^ The BARNABA software^77^ is used to calculate the eRMSD values of the RNA conformations with respect to the structure obtained after the energy minimization of the crystal structure. R_*ee*_ is the distance between the C4′ atoms of the terminal residues U67 and A74, and is computed using the post processing tools provided by GROMACS. The coordination number between different metal ions and a set of RNA atoms from the nucleobase and sugar fof the four key nucleotides, C68, C69, A72 and G73 is calculated using the PLUMED library^78,79^ (version 2.5.2). Details of the parameters used are provided in the supplementary information (SI).

### Spatial distribution of metal ions around RNA

To quantify the condensation of positively charged metal ions onto the negatively charged RNA, we calculated the local ion concentration (*c\**) around the different polar atoms present in the RNA (O atoms in phosphate, O atoms in sugar and, O and N atoms in nucleobases). Following earlier work,^26^ *c\** in molar units is defined as,

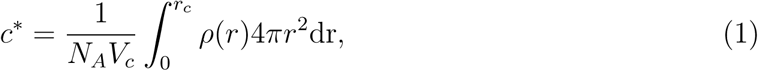

where *ρ*(*r*) is the number density of a specific type of ions at the distance *r* from a specific RNA atom, *V*_*c*_ is the spherical volume of radius *r*_*c*_, and *N*_*A*_ is the Avogadro’s number. The cutoff distance, *r*_*c*_ is set to 0.73 nm, which is the Bjerrum length of water at *T* = 300 K.

To study the distribution of different metal ions around the RNA residues in different basins of FES, we computed the spatial density maps using the volmap plugin of the VMD software.^80^ Initially, all the frames belonging to a particular basin are aligned to a representative RNA structure of the basin, which is randomly selected. The alignment is performed using the Kabsch method^81^ implemented in VMD. To construct the spatial density maps, the metal ions are treated as spheres of radii equal to their respective van der Waals radii and a grid of size 0.01 nm^3^ is constructed to divide the space around the RNA residues. A grid point is considered to be occupied and assigned a value equal to 1, if the sphere corresponding to a specific metal ion lies on the grid. Otherwise it takes a value equal to 0. The value at a grid point is averaged over all frames that belong to a specific basin. RNA residues, around which the occupancy of metal ions are found to be the highest in the spatial density maps, are further probed using the pair distribution functions, *g*(*r*). The *g*(*r*) between selected RNA residues and specific metal ions are calculated using VMD software with a distance cutoff of 2 nm and a grid spacing of 0.01 nm. Packages in VMD are also used for structural superposition^82^ and rendering of images.^83^

### Computing the TSE

To identify the TS structures that connect the folded and unfolded states, we have initially identified fragments of the simulation trajectory that either (a) leave the unfolded basin and enter the folded basin without returning to the unfolded basin or visiting any other basin, or (b) leave the folded basin and enter the unfolded basin without returning to the folded basin or visiting any other basin. Each such transition path, *TP*, detected in our trajectories (shown in the SI) is labeled as ‘reactive’ (Figure S2,S3). We have used the theory proposed by Hummer^84,85^ to identify the TSs in tetraloop folding. According to this theory, TSs are those points in the configuration space that have the highest probability of equilibrium trajectories passing through them are reactive. ^84^ Given that the system is in configuration *r*, the probability of being on a *TP* is given by

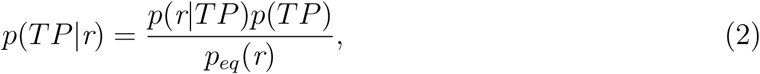

where *p*(*TP*) is the probability of a frame of the equilibrium trajectory to be on a TP, *p*_*eq*_(*r*) is the probability of a frame of the equilibrium trajectory to have the configuration *r*, and *p*(*r*|*TP*) is the probability of a frame of the equilibrium trajectory to have the configuration *r*, given that it already belongs to a TP. The configuration *r* is defined using the two CVs eRMSD and R_*ee*_. There are 46 and 97 tetraloop conformations in the simulation trajectories corresponding to [Mg^2+^] = 0 mM and 65 mM, respectively that have the maximum *p*(*TP*|*r*) value and are putative TS structures.

To verify that these conformations are TS structures, we further performed *P*_*fold*_ analysis.^86^ In the *P*_*fold*_ analysis, with each putative structure as an initial starting conformation (RNA + ions + water), we spawned 100 simulation trajectories each with a different set of velocities for a time period of 1.6 ns to probe whether they land in the folded or unfolded basins with approximately equal probabilities. For each starting structure, *P*_*fold*_ is defined as the fraction of these trajectories that land in the folded state. We labeled a putative structure as a TS, if the *P*_*fold*_ computed from 100 independent simulation runs satisfies 0.4 *≤ P*_*fold*_ *≤* 0.6. Representative trajectories are shown in the SI (Figure S4C,D). Using this procedure, we have identified four conformations as the TS structures for each of the [Mg^2+^] = 0 mM and 65 mM systems.

## Results and Discussion

### Folded Structure in the Simulations Deviates from the Crystal Structure

Two independent simulations of the L4 tetraloop hairpin motif performed at *T* = 300 K and [Mg^2+^] = 0 mM for a cumulative time of ≈ 20 *µ*s long show multiple transitions between the folded and unfolded conformations (Figure S1A,B). A total of 8 folding events are observed in both the trajectories. Analyses of the folding events using CN_*K*_ reveal active involvement of K^+^ in L4 tetraloop folding. The folded conformations are short lived with an average lifetime of ≈ 2.5 ns. We also performed two independent simulations of the L4 tetraloop at *T* = 300 K and [Mg^2+^] = 65 mM for a cumulative time of ≈ 20 *µ*s. We observed 10 folding events in these trajectories (Figure S1C,D). In the presence of Mg^2+^, the average life time of the folded state increased to ≈ 8.5 ns indicating that Mg^2+^ stabilize the folded state compared to K^+^.

There are structural differences between the folded conformation of the L4 tetraloop obtained from the MD simulations and crystal structure (Figure S5). In the crystal structure,^49^ the geometry of the L4 teraloop at the termini is influenced by long range interactions, which are absent in the isolated system modeled in MD simulations (Figure S5A,B). For example, the helix of the L4 hairpin is coaxially stacked with the P4 helix (Figure S5C,D), and also A71 and A72 residues from the loop of the L4 hairpin are involved in hydrogen bonding interactions with the P5 stem and L5 loop, respectively (Figure S5E). Although the deviation between the crystal structure and the folded structure in the MD simulations is significant at the termini and at the orientation of the C69 residue (Figure S5A,B), the consecutive stacking interactions between the bases of A70, A71 and A72 remain relatively unperturbed (Figure S5F).

### Site Specific Binding of a Metal Ion is Indispensable for the Folding of L4 Tetraloop

We constructed the FES of L4 tetraloop folding to probe the role of metal ions in the folding mechanism (Figure 2). We projected the FES onto the following CVs: eRMSD, CN_*K*_ and CN_*Mg*_, where CN_*K*_ and CN_*Mg*_ are the coordination numbers of K^+^ and Mg^2+^ with respect to the atoms of four nucleosides C68, C69, A72 and G73. Note that C69 and A72 form the loop closing pair, which is followed by the first base pair of the helix composed of C68 and G73 (Figure 1). Therefore, when a metal ion gets trapped inside the RNA loop, its corresponding value of coordination number increases. Low and high values of eRMSD correspond to folded and unfolded conformations, respectively.

**Figure 2:**
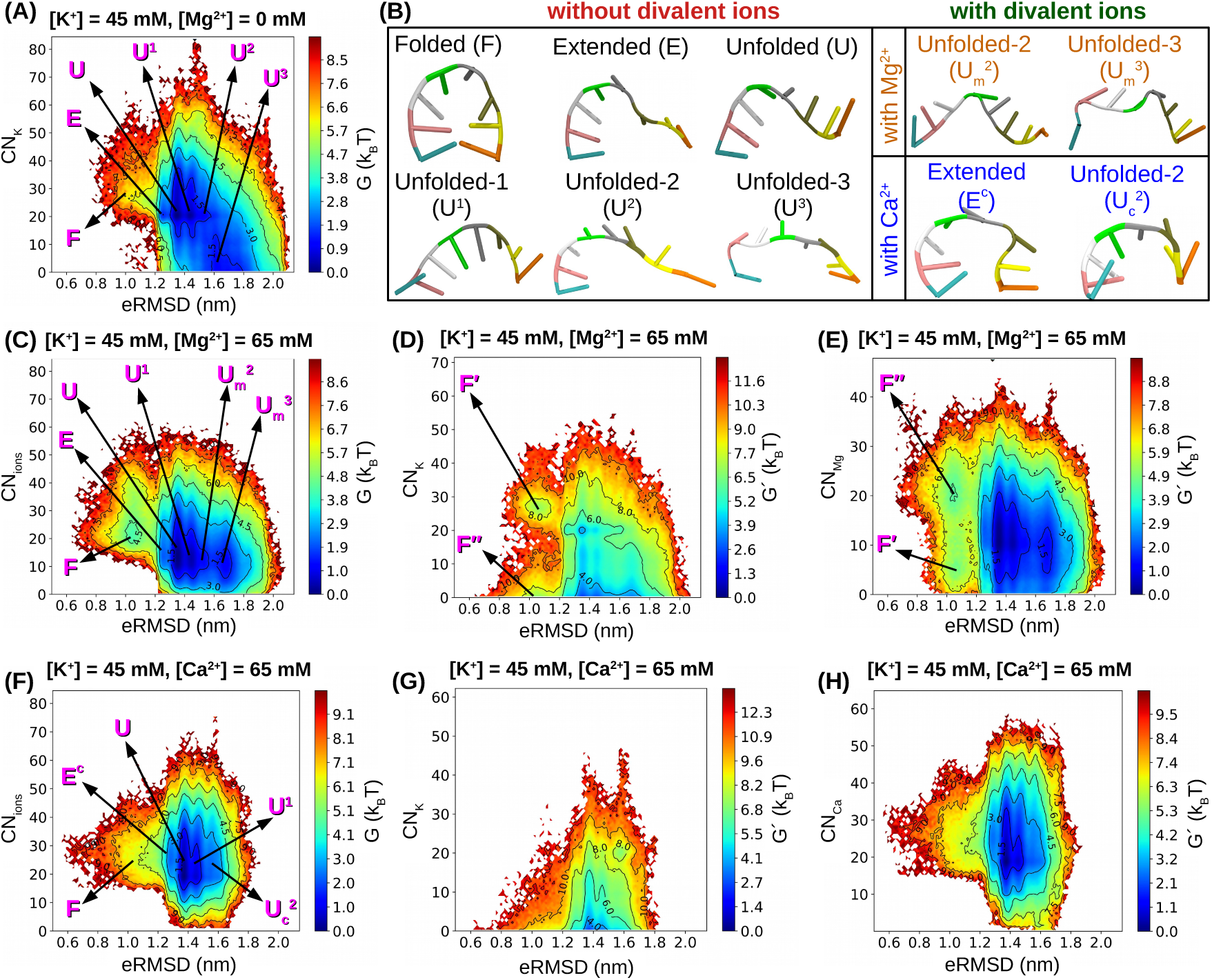
(A) FES of the L4 tetraloop projected onto CVs, eRMSD and CN_*K*_ at [Mg^2+^] = 0 mM. Important basins in the FES are highlighted. (B) Representative structure of the RNA from different basins are shown in cartoon representation. (C) FES projected onto eRMSD and CN_*ions*_ (= CN_*K*_ + CN_*Mg*_) at [Mg^2+^] = 65 mM. In a solution containing K^+^ and Mg^2+^, FES is further projected onto eRMSD, CN_*K*_ and CN_*Mg*_. (D) FES constructed using 3 CVs is shown in two dimensions, eRMSD and CN_*K*_, after integrating out the third dimension, CN_*Mg*_. (E) Similarly, FES is projected onto eRMSD and CN_*Mg*_ after integrating out CN_*K*_. In these projections, instead of a single folded basin ‘F’, two folded basins F′ and F″ are visible, which are due to the specific binding of K^+^ and Mg^2+^, respectively. (F) FES projected onto eRMSD and CN_*ions*_ (= CN_*K*_ + CN_*Ca*_) at [Ca^2+^] = 65 mM. In a solution containing K^+^ and Ca^2+^, FES is further projected onto eRMSD, CN_*K*_ and CN_*Ca*_. (G) FES is shown in 2 dimensions, eRMSD and CN_*K*_, after integrating out CN_*Ca*_. (H) Similarly, FES is projected onto eRMSD and CN_*Ca*_ after integrating out CN_*K*_. In a solution containing K^+^ and Ca^2+^, only Ca^2+^ contributed to L4 tetraloop folding.

The FES for both [Mg^2+^] = 0 mM (projected onto eRMSD and CN_*K*_) and 65 mM (projected onto eRMSD and CN_*ions*_ (= CN_*K*_ + CN_*Mg*_)) solutions shows 3 distinct basins, which correspond to (i) the folded state (basin ‘F’ at eRMSD ≈ 1.04 nm), (ii) a semi-folded state with extended end-to-end distance (basin ‘E’ at eRMSD ≈ 1.25 nm) and (iii) a broad unfolded basin (Figure 2A,C). The unfolded basin can be further classified into 4 basins. RNA conformations that populate the basin ‘U’ centered around eRMSD ≈ 1.35 nm, have all the eight bases stacked consecutively (Figure S5G). The overall backbone orientation resembles that of a single stranded helix, as reported for homopolymeric RNAs.^87^ The other three basins correspond to different unfolded conformations of the the RNA and are centered around eRMSD ≈ 1.44 (basin U^1^), ≈ 1.53 (basin U^2^) and ≈ 1.67 nm (basin U^3^), respectively (Figure 2A,C). In the presence of Mg^2+^, the basins located at higher values of eRMSD (U^2^ & U^3^) get populated with RNA conformations that have noticeable variation in their backbone orientation (Figure 2B).

The FES shows that both Mg^2+^ and K^+^ ions contribute to folding. When [Mg^2+^] = 0 mM, the folded basin (F) at CN_*K*_ ≈ 27.5 and eRMSD ≈ 1.04 nm is narrow and shallow (Figure 2A). The absence of a basin close to CN_*K*_ ≈ 0 and low eRMSD values suggests that binding of K^+^ within the region surrounded by C68-C69-A72-G73 residues is indispensable for the folding of L4 tetraloop. In the presence of Mg^2+^, the FES projected onto eRMSD and CN_*ions*_ (= CN_*K*_ + CN_*Mg*_) shows a broad and deep folded basin with CN_*ions*_ 21 and eRMSD ≈ 1.04 nm (Figure 2C). Even in this case, the absence of a basin close to CN_*ions*_ ≈ 0 and low eRMSD values indicate that the folding of L4 tetraloop is driven by the specific binding of a metal ion (either Mg^2+^ or K^+^) inside the loop. Representative structures of the folded state with the metal ion (K^+^/Mg^2+^) bound at the center of the loop are illustrated in Figure S6.

### Mg^2+^ Stabilizes the Folded Structure Compared to Other Metal Ions

To contrast the L4 tetraloop folding in the presence of Mg^2+^ and K^+^, we compared the FES at [Mg^2+^] = 0 mM and 65 mM (Figure 2A,C). The folded state becomes more stable when Mg^2+^ is also present in the solution in addition to K^+^ (Figure 2C). This is clear from the FES projected onto a single CV, eRMSD (Figure S7). The free energy difference between the folded and unfolded states is estimated^34^ as Δ*G*_*F U*_ = −*k*_B_*T* [log(∑_*i,F*_ *P* (**s**_*i*_)) − log(∑_*i,U*_ *P* (**s**_*i*_))], where F and U correspond to the folded and unfolded basins, respectively (Figure 2A,C). ΔG_*F U*_ for the L4 tetraloop decreases from 14.6 kJ/mol to 10.6 kJ/mol on increasing [Mg^2+^] from 0 mM to 65 mM. This value is significantly less than the ΔG_*F U*_ (≈ 16.1 kJ/mol) of a standard GNRA tetraloop at the same temperature in the presence of only monovalent counterions.^34,35^ However, contribution of K^+^ in stabilizing the L4 tetraloop folded state when both K^+^ and Mg^2+^ are present in the solution remains ambiguous from the FES shown in Figure 2C.

UV melting experiments^88^ and computational studies^89^ have established that when monovalent ions are also present in the solution along with Mg^2+^, they compete in stabilizing RNA tertiary structures. At low Mg^2+^ concentration ([Mg^2+^] ≈ 0), RNA’s stability primarily depends on the monovalent ion concentration, whereas at higher [Mg^2+^] it becomes independent of the concentration of the monovalent ion. To probe the contribution of K^+^ and Mg^2+^ to L4 tetraloop folding when both K^+^ and Mg^2+^ are present in the solution, we projected the FES onto three CVs, eRMSD, CN_*K*_ and CN_*Mg*_. The FES shows that even in the mixture of K^+^ and Mg^2+^, both ions contribute to the folding of the tetraloop (Figure 2D,E). The two folded basins, F′ and F″, observed in the FES are due to the specific binding of K^+^ and Mg^2+^, respectively in the center of the hairpin motiff formed in the folded state. However, the folded basin F″ is deeper compared to the folded basin F′ showing that RNAs fold efficiently in the presence of Mg^2+^ at the center of the loop compared to the K^+^.

We further studied whether other divalent ions such as Ca^2+^ are as efficient as Mg^2+^ in stabilizing the folded state of L4 tetraloop. We performed two independent simulations of the L4 tetraloop at *T* = 300 K and [Ca^2+^] = 65 mM for a cumulative time of ≈ 20 *µ*s long (Figure S1E,F). A total of 3 folding events are observed in both the trajectories. The FES projected onto eRMSD and CN_*ions*_ (= CN_*K*_ + CN_*Ca*_) does not show any deep folded basin as observed for 65 mM Mg^2+^ solution (Figure 2F). This shows that the increase in the folded state stability in the presence of Mg^2+^ is not just due to the abundance of cations present in the medium. Also the RNA conformations that populate the semi-folded extended basin labeled ‘E’ (eRMSD ≈ 1.25 nm) is distinctly different from the conformations that populate the same basin in [Mg^2+^] = 0 mM and [Mg^2+^] = 65 mM systems (Figure 2B). Absence of a basin close to CN_*ions*_ ≈ 0 and low eRMSD suggests that even in this case, the folding of L4 tetraloop is driven by the specific binding of metal ions (either Ca^2+^ or K^+^) at the center of the loop. However, the FES projected onto eRMSD, CN_*K*_ and CN_*Ca*_ has no basin at eRMSD *<* 1.2 nm for either at a high value of CN_*K*_ (Figure 2G) or at CN_*Ca*_ ≈ 0 (Figure 2H). This implies that in the presence of Ca^2+^, the monovalent K^+^ did not contribute towards folding and we explore the reasons below.

### Ca^2+^ and Mg^2+^ Condense More Efficiently on the RNA Compared to K^+^

Monovalent metal ions like K^+^ can actively participate in stabilizing RNA tertiary structures,^90^ selectively chelate to specific sites on RNA (e.g. AA platform,^27^ G-quadruplex,^91^ etc.) and even modulate the folding pathways of RNA.^20^ Yet K^+^ show limited participation in modulating the folding of L4 tetraloop, especially when the solution contains a mixture of divalent and monovalent ions. This can be rationalized from the difference in the ion condensation patterns around the RNA (Figure 3). It was shown earlier that in a mixture of divalent and monovalent ions, divalent ions are preferred over monovalent ions for condensation around the negatively charged RNA, due to their high charge to volume ratio. ^26^ We also observe that in the absence of divalent ions, the K^+^ condense around both backbone (sugar-phosphate) and nucleobase atoms (blue line in Figure 3). However, in the presence of divalent ions, K^+^ concentration around the RNA atoms drops drastically (magenta and cyan lines in Figure 3). In the mixture of K^+^ and Ca^2+^, the K^+^ have *c\** ≈ 0 at all the polar sites on RNA (magenta line in Figure 3), as only the Ca^2+^ condense around the polar sites of RNA (*c\** 0.5 for most of the sites, black line in Figure 3). However, in the mixture of K^+^ and Mg^2+^, a noticeable concentration of K^+^ is observed around the RNA atoms with relatively higher *c\** values around the nucleobase atoms present at the Hoogsteen edge of the purine residues (Figure 3D-H). Therefore in a mixture of K^+^ and Mg^2+^ both contribute towards folding with Mg^2+^ being the major contributor (Figure 3D,E). The important point to note here is that all the three metal ions (K^+^, Mg^2+^ and Ca^2+^) prefer to condense around the Hoogsteen edge of A72/G73. Therefore, the ion binding site (as present in the crystal structure) cannot discriminate between different metal ions.

**Figure 3:**
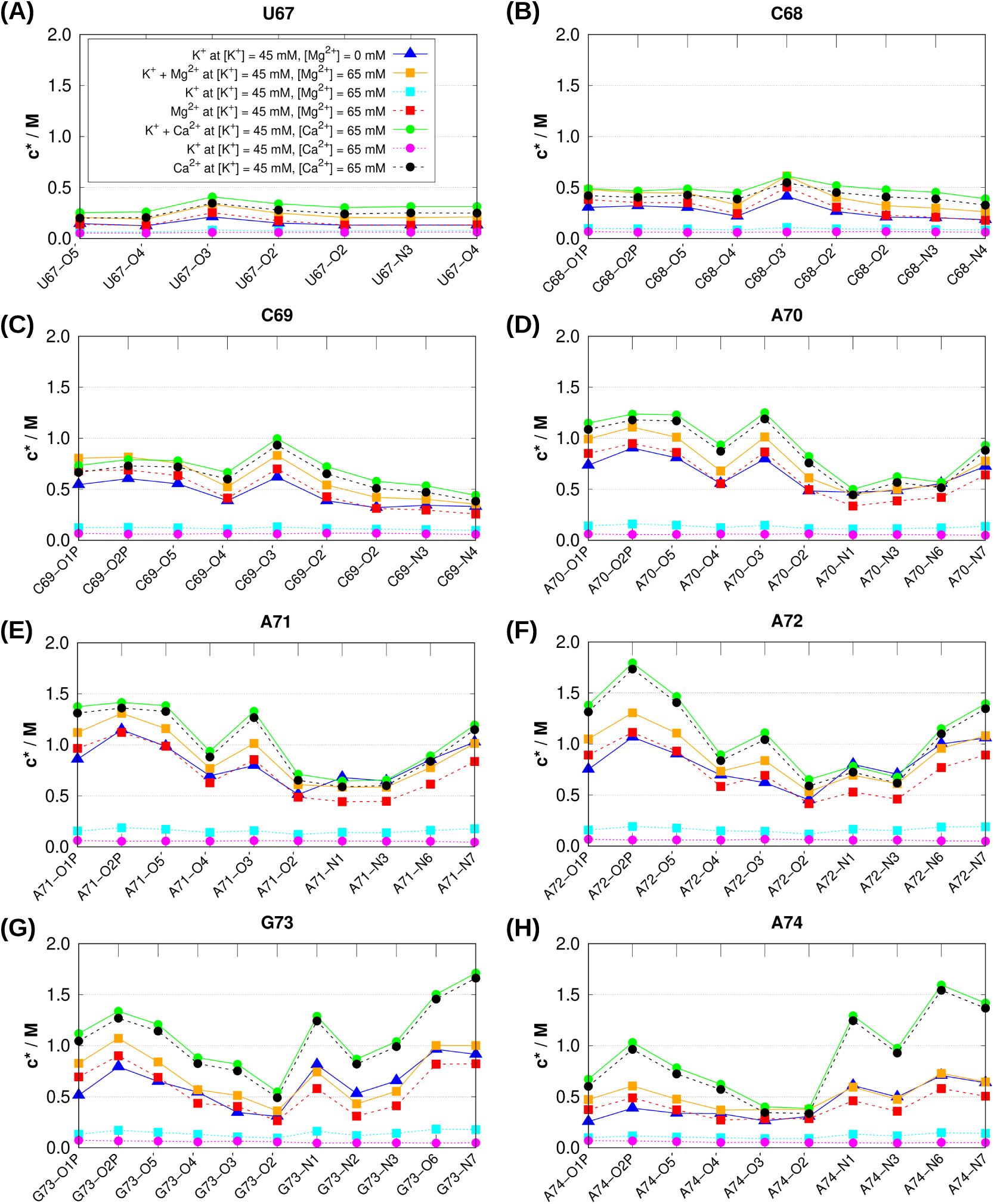
Fingerprint of ion coordination expressed as local ion concentrations (*c\**) at different polar sites located on the eight RNA residues (A) U67, (B) C68, (C) C69, (D) A70, (E) A71, (F) A72, (G) G73 and (H) A74. Different binding sites on the RNA are annotated following the PDB nomenclature.

This observation is also in agreement with the experiments,^49,92,93^ which have shown that Mg^2+^ sensing M-box riboswitch from which the RNA fragment is extracted is not very selective for binding Mg^2+^ unlike other metal sensing riboswitches. This riboswitch is shown to compact and fold in the presence of other divalent metal ions such as Mn^2+^. Crystals of M-box riboswitch grown in the presence of Mn^2+^ show that the binding pockets of Mg^2+^ are occupied by Mn^2+^.^92,93^ Since the Mg^2+^ sensing M-box riboswitch has small affinity and selectivity of metal ions it is hypothesised that this riboswitch probably functions by binding to the most prevalent Mg^2+^ ion present in the cell.

### Specific Site Bound Metal Ion is Intrinsic Component of the TS

Although, the L4 tetraloop folds into the shape of a conventional GNRA tetraloop in the presence of Mg^2+^, its folding thermodynamics is significantly different from that of a conventional GNRA tetraloop. The rate limiting step in the unfolding of GNRA tetraloops is the breaking of the loop closing G:A *trans*S:H base pair^94^ stabilized by two strong inter-base hydrogen bonds.^95^ Whereas the crystal structure of L4 tetraloop suggests that stability of the loop closing C:A pair and hence, the sequential adenine-adenine stacks primarily depend on the presence of hydrated Mg^2+^ inside the loop (Figure 1). To probe at what stage of folding, the metal ion binds to the RNA and how it selects the specific site for binding, we further identified the TS connecting the folded and unfolded basins.

We have used the theory proposed by Hummer^84,85^ to identify the TSE using the CVs, eRMSD and R_*ee*_. The FESs projected onto eRMSD and R_*ee*_ show that R_*ee*_ can also resolve the folded state and various unfolded states for both [Mg^2+^] = 0 mM and [Mg^2+^] = 65 mM (Figure S8). In the previous studies of GAGA and UUCG type tetraloop folding, eRMSD and R_*ee*_ were also used as CVs to construct the folding FES.^34,35^ To identify TSE, we analyzed the paths that connect the unfolded and folded basins of the L4 tetraloop in the simulation trajectories. In each path, the RNA fragment before transitioning to the folded basin hops between different unfolded basins and the extended basin. The final transition to the folded basin F takes place from either the unfolded basin U or from the extended basin E. In the absence of Mg^2+^, 20% of the transitions to the folded state took place from the extended basin. Whereas, in the presence of Mg^2+^ it reduced to only 8%. Therefore we restricted our TS search to only the paths connecting the unfolded U and folded F basins (Figure S2,S3).

TSE analysis carried out for [Mg^2+^] = 0 mM and 65 mM systems suggests that for both the systems, the value of *p*(*TP*|*r*) is maximum at the same configuration *r*, which is located at eRMSD ≈ 1.2 nm and R_*ee*_ ≈ 1.8 nm on the FES (Figure S4A,B). Further, *P*_*fold*_ analysis of the RNA conformations (along with the ion and water molecules) that have maximum value of *p*(*TP*|*r*) helped in identifying the role of ions and water in the TS structures. The overall geometry of the TS structures are similar when folding is driven by the specific binding of K^+^ or Mg^2+^ inside the tetraloop (Figure 4). The TS is primarily composed of an unfolded helix and a nearly folded loop with a site bound metal ion. However, the interactions between RNA and the metal ions change significantly when Mg^2+^ is added to the solution.

**Figure 4:**
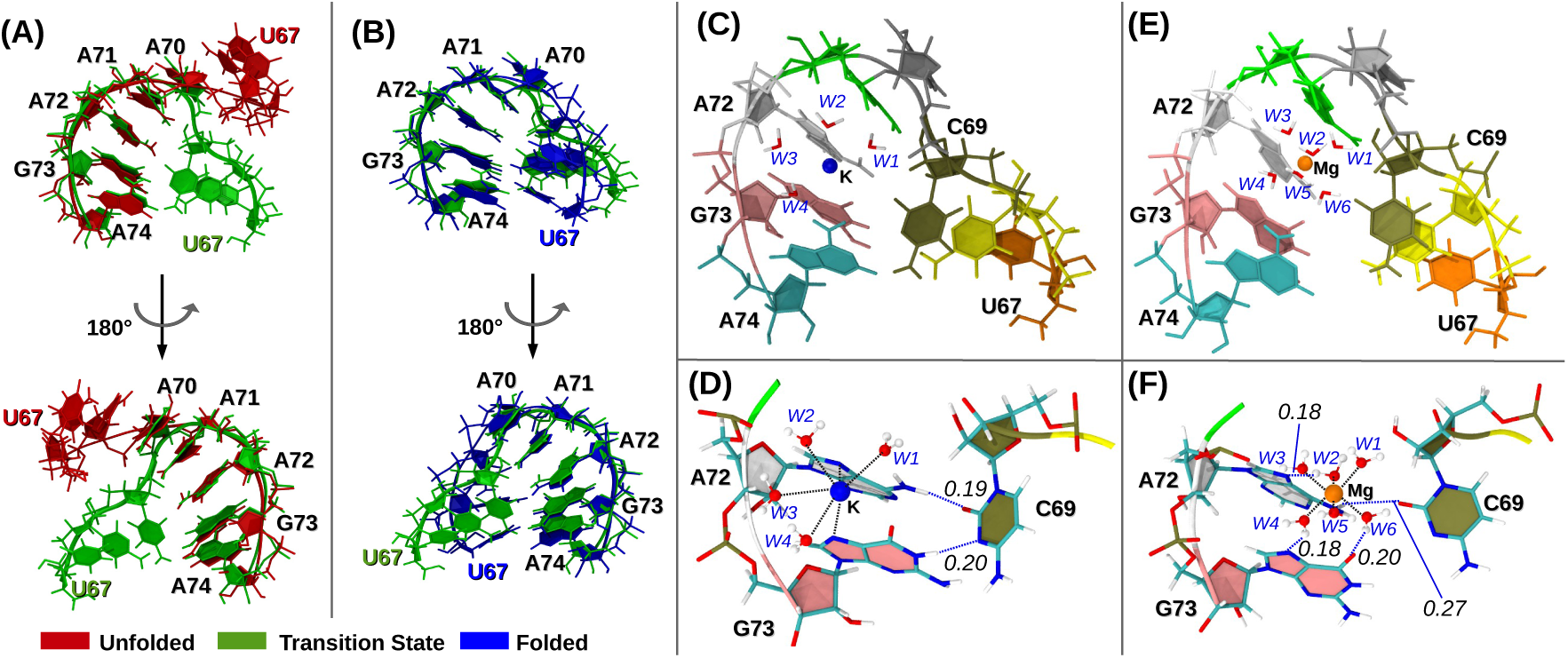
Representative structure from the TSE (green) is structurally aligned with a representative structure from the (A) folded basin (blue) and (B) unfolded basin (red). The alignment is performed with respect to only the A70-A74 residues. (C) Representative structure of the TSE with a K^+^ bound at the Hoogsteen edge of A72 and G73. The K^+^ is partially dehydrated and four water molecules present in its first coordination shell are labeled W1 - W4. (D) Details of the K^+^-RNA interactions are further illustrated. The K^+^ directly interacts with the N7 atoms of A72 and G73 and with four water molecules. These contacts are shown in black broken line. C69 forms two inter-base hydrogen bonds (shown in blue broken lines), one between O2 of C69 and N6 of A72, and the other between N3 of C69 and N1 of G73. (E) Representative structure from the TSE shows a hydrated Mg^2+^ bound at the Hoogsteen edge of A72 and G73. The six water molecules in the first coordination shell of the bound Mg^2+^ are labeled W1 - W6. (F) Details of the Mg^2+^-RNA interactions are highlighted. The W2, W4 and W6 water molecules form hydrogen bonds with the N7 atom of A72, N7 atom of G73 and O6 atom of G73, respectively. In addition there is an inter-base hydrogen bond between O2 of C69 and N6 of A72. These hydrogen bonds are shown in blue broken lines. Interaction between Mg^2+^ and water molecules are shown in black broken lines. For each hydrogen bond, the corresponding distance between the hydrogen and the acceptor is reported in nm. Metal ions are shown as spheres with radii equal to 30% of their respective van der Waals radius.

### Hydrated Mg^2+^ Forms Key Interactions in the TS that Drive the Formation of the Loop

Structural superposition of (a) unfolded and TS structures (Figure 4A) and, (b) folded and TS structures (Figure 4B) collectively suggest that for the formation of the folded hairpin motif from the single stranded helix conformation in the unfolded state will require breaking of the base-base stacking interactions between the consecutive C69 and A70 residues at the 5′ end. This is followed by the formation of the A-form helix involving C68:G73 and U67:A74 base pairs. In DNA double helix, base-stacking is the main stabilizing factor.^96^ Similarly in single stranded RNA helix, the strong base-stacking interactions^97^ are important for structural stabilization. Therefore, during the L4 tetraloop folding, after the breakdown in the consecutive base-stacking interactions at the 5′ end, the resulting alternate geometry has to be stabilized by other competing noncovalent interactions leading to folding. Structures in the TSE reveal that it involves a network of hydrogen bonding interactions that connect C69 to the stacked A72/G73 residues (Figure 4C,E). This network of hydrogen bonding interactions are further supported by metal ion coordination.

In the case of K^+^ driven folding, a partially dehydrated K^+^ is observed to interact directly with the Hoogsteen edges (N7 positions) of A72 and G73 (Figure 4D). Whereas, in the case of Mg^2+^ driven folding (Figure 4F), the water molecules present in the coordination shell of the Mg^2+^ interact with the Hoogsteen edges of both A72 (N7 position) and G73 (both N7 and O6 positions). The resulting electrostatic interactions between this cationic environment and the electronegative sites of the WC edge of C69 (O2 and N3) are crucial for the stabilization of the loop in the TS. Compared to other metal ions, Mg^2+^ has an extremely high charge to volume ratio due to its small ionic radius (Table S1), which makes it a preferable candidate for mediation of these electrostatic interactions.

Quantum chemical calculations suggest that metal ion coordination at the Hoogsteen edge of purines modulate the charge distribution and hence the hydrogen bonding potential of the respective WC edges.^98–100^ Generally, on metal ion binding at the Hoogsteen edge of a purine, the hydrogen bonding potential of the hydrogen bond donors (N6 in adenine, N1 and N2 in guanine) and acceptors (N1 in adenine, O6 in guanine) increases and decreases, respectively. However, in classical MD simulations with non-polarizable force fields we cannot capture such effects.

### A-form Helix Formation Requires Rearrangement of Mg^2+^ Mediated Interactions

Comparison of the structures in the TSE and folded state shows that C69 and A72 interact with each other in *trans* orientation in the folded state (Figure 4B,S6B,D), whereas in the TSE they interact in *cis* orientation (Figure 4D,F). Therefore for the L4 tetraloop to fold, the C69 residue has to rearrange itself to interact with A72 in *cis* conformation. This requires breaking of the hydrogen bond between O2 of C69 and N6 of A72 present in the TS (Figure 4D,F) and formation of new hydrogen bonds between O2 of C69 and water molecules present at the first coordination shell of K^+^/Mg^2+^ (Figure S6B,D).

For K^+^ driven folding, this rearrangement results in the formation of new hydrogen bond between one K^+^ coordinated water and O6 position of G73 in the folded state (Figure S6B). Also, N6 of A72 forms a hydrogen bond with the sugar oxygen O2′ of C69 (Figure S6B). For Mg^2+^ driven folding (Figure S6D) the consequence of such rearrangement is reflected in elongation of the hydrogen bonds that connect the hydrated Mg^2+^ to the Hoogsteen edges of A72 and G73. This shows that the K^+^/Mg^2+^ coordinated water molecules are crucial for rearrangement of C69 residue and subsequent formation of the A-form helix. Divalent cations like Ca^2+^ that lack the rigid octahedral orientation of the inner-shell water molecules^101^ cannot efficiently facilitate the formation of L4 tetraloop hairpin motif. Hence, the role of Mg^2+^ in the folding of L4 hairpin is very specific and other divalent cations are not as efficient in creating the same effect.

### Spatial Organization of Electronegative Nucleobase Atoms Drive the Site-Specific Binding of Mg^2+^

Mg^2+^ binds to specific sites in RNA and controls its folding.^42^ The specific binding was proposed to be driven by spatial arrangement of negatively charged phosphate groups in the folded RNA molecule and subsequent development of binding pockets with high anionic potential. However, using a coarse-grained model of *Azoarcus* ribozyme and MD simulations it was recently demonstrated that site specific binding of Mg^2+^ to RNA occurs even when the RNA is unfolded and the tertiary interactions that define the folded conformation are absent.^26^

In our simulations, we also observe that Mg^2+^ binds to the Hoogsteen edge of A72 even when the L4 tetraloop is unfolded. The spatial density map of the metal ions around the RNA conformations in the unfolded state show that occupancy of the metal ions is maximum near the Hoogsteen edge of A72 (Figure 5A-5D). To probe the interaction between metal ions and the Hoogsteen edge of A72, we computed the pair distribution function (*g*(*r*)) between different metal ions and the RNA fragment (Figure 5E). Similar to the trends observed in Figure 3, the *g*(*r*) peak height for Mg^2+^ is significantly higher than that for K^+^ in [Mg^2+^] = 65 mM solution. It is easier for the metal ions with smaller size such as Mg^2+^ to reach specific RNA binding pockets with high electronegative potentials, as proposed in earlier reports.^26,42^ Consistent with the earlier report^102^ we also observe that the position of the first peak in the *g*(*r*) for K^+^ is at a shorter distance (*r* ≈ 0.2 nm) compared to that for Mg^2+^ (*r* ≈ 0.5 nm). This indicates that K^+^ directly interacts with the RNA, whereas Mg^2+^ interacts via the water molecules in its first coordination shell (diameter of water molecule is ≈ 0.28 nm). Interestingly, the occupancy of metal ions is found to be maximum near the Hoogsteen edge of A72 even for RNA conformations in the folded basin (Figure 5). We probed further to understand the reason for consistent electrostatic potential around the Hoogsteen edge of A72, which facilitates the binding of metal ions throughout the L4 tetraloop folding process. Superposition of folded, unfolded and TS structures with respect to each other reveal that, throughout the folding process five consecutive purine residues at the 3′ end (A70-A74) remain stacked with each other (Figure 4A-B). This brings the Hoogsteen edges of the five purines in a specific order where the Hoogsteen edge of A72 remains at the center. It is known that hydrogen bond acceptor sites of the Hoogsteen edges of purines (N7 in adenine and N7 and O6 in guanine) are strongly electronegative^103^ and N7 of guanine has the highest proton affinity out of all the nucleobase atoms.^104^ Therefore, we argue that in addition to the negatively charged backbone phosphate groups, spatial organization of electronegative nucleobase atoms critically contribute towards construction of the unique electrostatic potential that drives metal ion binding at specific sites. To find out whether it is a general rule or specific to this L4 tetraloop hairpin motif, requires further investigation.

**Figure 5:**
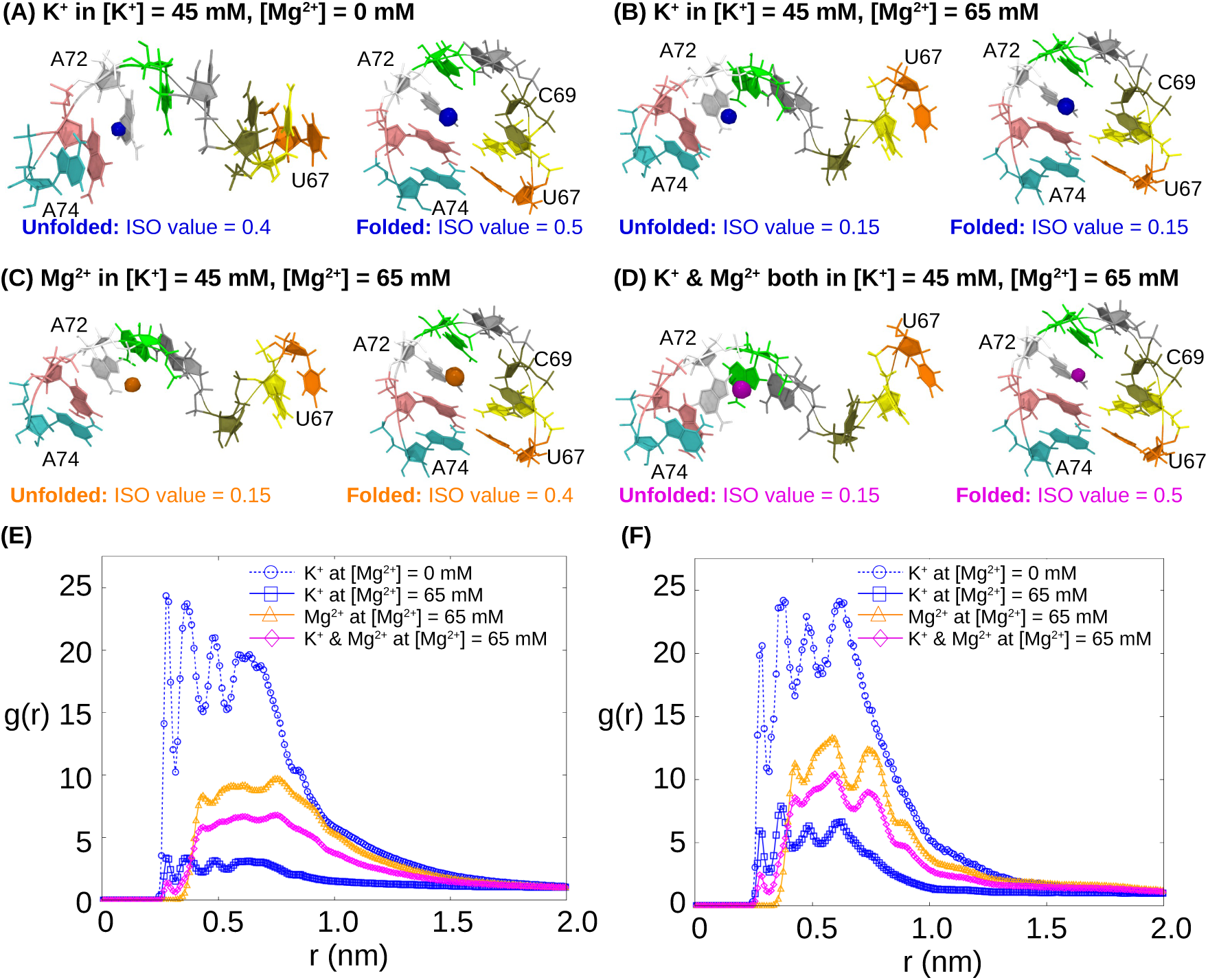
(A) Spatial density map of K^+^ around the unfolded and folded states of RNA are shown as blue isosurfaces at [Mg^2+^] = 0 mM. Respective ISO value of a isosurface indicates the fraction of time for which the area under the surface is occupied by K^+^. Similarly, at [Mg^2+^] = 65 mM, spatial density maps for (B) K^+^, (C) Mg^2+^ and (D) both K^+^ & Mg^2+^ around the unfolded and folded states of RNA are shown as isosurfaces in blue, orange and magenta, respectively. ISO values for each isosurfaces are mentioned below the respective illustrations. Maximum observed ISO values for each case are given in Table S2. (E) Pair distribution function (*g*(*r*)) between 2 groups of atoms for the RNA unfolded state. The first group contains atoms of A72 and G73 nucleotides of RNA, and the second group contains metal ions present in the solution. *r* is the distance between a pair of particles in the 2 groups. (F) *g*(*r*) between 2 groups of atoms for the RNA folded state. The first group contains atoms of C69 and A72 nucleotides of the RNA and the second group contains different metal ions present in the solution.

The results discussed above collectively suggest that binding of metal ions at specific site on RNA (hoogsteen edge of A72/G73) is indispensable for the folding of L4 tetraloop. Folding efficiency becomes maximum when the bound ion is Mg^2+^. Such selective preference towards Mg^2+^ is determined by its involvement in the TS. In fact, hydrated Mg^2+^ are intrinsic components of the TS. Binding of hydrated Mg^2+^ at the Hoogsteen edge of A72/G73 provides the electrostatic stabilization necessary for the formation of the loop part of the motif. For the subsequent formation of the helix part of the motif, the Mg^2+^ coordinated water molecules actively mediate the rearrangement of the necessary inter-nucleotide interactions. Other divalent ions like Ca^2+^ that lack a rigid octahedral arrangement of their coordinated water molecules fail to facilitate these rearrangements efficiently. On the other hand, monovalent ions like K^+^ has to undergo partial dehydration to facilitate folding. This is energetically costly^105,106^ and the subsequent folded state is also unstable. We further observe that the specific electronegative potential required for the binding of Mg^2+^ is provided by the spatial organization of the electronegative nucleobase atoms of selected nucleotides, in addition to the negatively charged backbone. Therefore, the CVs designed on the basis of coordination number between metal ions and atoms of selected nucleosides (nucleobase + sugar) successfully provide a comprehensive description of the free energy landscape.

## Conclusions

Several aspects of RNA folding remains elusive despite extensive efforts.^107^ Elucidating the exclusive contribution of metal ions in the formation of globular RNA structures is one of the major challenges.^108^ Out of various metal ions, Mg^2+^ has the most versatile contribution in RNA’s folding. Recent studies have unraveled active participation of Mg^2+^ in modulating RNA’s catalytic activity,^109^ conformational flexibility,^110^ and stability of folding intermediates.^111^ In this paper we show that a site bound hydrated Mg^2+^ is an intrinsic component of the TS of a unique RNA hairpin motif. Similar to the folding of the P4-P6 RNA domain, ^112^ the observed TS is an early TS which lacks majority of its native contacts. Formation of these missing native contacts takes place through water mediated interactions promoted by the water molecules present in the first coordination shell of the chelated Mg^2+^. Mg^2+^ co-ordinated waters are known to form strong hydrogen bonds in RNA.^113^ The L4 tetraloop hairpin motif studied here utilizes such strong hydrogen bonding interactions to connect the distant nucleotides and stabilize the folded state. This is an important addition to the long standing discussion^114,115^ on the importance of water in RNA folding.

The larger goal of our study was to understand how large RNAs, like metal sensing riboswitches,^48^ selectively recognize specific metal ions. Conventionally anionic phosphate groups are treated as building blocks of metal binding pockets in RNA.^36^ However, the metal ion binding site present in the L4 tetraloop is composed of electronegative nucleobase atoms. This is not surprising as structural bioinformatics based analysis of available RNA crystal structures have revealed that out of the 13 known Mg^2+^ binding motifs in RNA, 8 have one or more nucleobase atoms present in the inner and/or outer shell of the Mg^2+^.^24^ Rather what is remarkable in this context is the fact that, this binding site cannot discriminate between different metal ions. Experimentally, it was shown that for Mg^2+^ sensing M-box riboswitch from which the RNA fragment is extracted is not very selective for binding Mg^2+^.^92^ Based on this it was argued that M-box riboswitches can sense Mg^2+^ because they are the most prevalent divalent ions *in-vivo*.^48,92^ However, we propose an alternative hyopthesis for the source of metal ion specificity in metallo-riboswitches. As we know, metal ion binding in riboswitches^116^ depends on the availability of a binding site with complementary electrostatic potential. In addition to that we propose the active involvement of the metal ion in the formation of the TS and stabilization of the folded state could also be an important factor. We further observe that, even for a small RNA fragment like the L4 tetraloop hairpin motif, the folding FES becomes multidimensional especially when multiple metal ions are present in the solution. We have identified relevant CVs, which will aid in computing the FES of large RNAs using enhanced sampling techniques. The L4 tetraloop (CAAA) folds in the shape of a conventional GNRA tetraloop, where two nucleobases (C69 and A72) connected by hydrated metal ions mimics the effect of an isosteric^117^ substitution of a noncanonical base pairs (i.e., the terminal G:A *trans*S:H pair present in a GNRA tetraloop). It is therefore important for the developers of RNA 2D^118^ and 3D^119^ structure prediction algorithms to take this check point into consideration. Interestingly, the nearest GNRA sequence of the L4 tetraloop is GAAA (Figure 1). Tertiary interaction between this GAAA tetraloop and its receptor motif has been used extensively as a model system in single molecule experiments to study various thermodynamic and kinetic aspects of RNA folding.^120^ They reveal a complex dependence of the RNA folding dynamics on metal ion concentrations. ^121,122^ Note that docking of the tetraloop with its receptor takes place via only the A-minor interactions involving the adenine residues of the tetralop, which are common in both GAAA and L4 tetraloops.^123^ Also it is demonstrated that these adenine residues remain spatially constrained throughout the folding of L4 tetraloop. Therefore, we suggest that studying the tetraloop-receptor interaction where the GAAA tetraloop is substituted by the metal sensitive L4 tetraloop may reveal further information regarding involvement of metal ions in RNA folding.

Scope of this work is limited by relatively small system size and application of a fixed charge based force field. Study of the complete aptamer domain of metal sensing riboswitches using polarized force fields can further validate our hypothesis. Nevertheless, to the best of our knowledge, this is the first study that elucidates the folding free energy surface of a RNA hairpin using unbiased all atom molecular dynamics simulations.

## Acknowledgement

This work was supported in part by a grant from the National Supercomputing Mission (NSM) through the grant MeitY/R&D/HPC/2(1)/2014. G.R. acknowledges the Alexander von Humboldt Foundation for support of his visit to the Max Planck Institute for Polymer Research in Mainz. A.H. acknowledges the financial support from the DBT-RA Program in Biotechnology and Life Sciences. S.K. acknowledges research fellowship from Indian Institute of Science-Bangalore. The computations are performed using the TUE and Cray XC40 clusters at IISc. The authors thank Joseph F Rudzinski and Arghya Dutta for critical reading of the manuscript.

## Supporting Information Available

Details of the coordination number calculation using PLUMED, list of parameters used for calculating the electrostatic potential surfaces in the PBEQ Solver server, Figure S1 – S8 and Table S1, S2. Figure S1: Time evaluation of eRMSD values in six equilibrium trajectories analyzed in this work. Figure S2: Illustration of transition paths in 0mM Mg^2+^ solution. Figure S3: Illustration of transition paths in 65mM Mg^2+^ solution. Figure S4: Results of the Hummer analysis performed to obtain the TSE. Figure S5: Differences between the crystal structure of the L4 tetraloop hairpin motif and its folded structure obtained in MD simulations. Figure S6: Details of metal-RNA interactions in the folded state. Figure S7: FES projected on single CV, eRMSD. Figure S8: FESs projected on two CVs, eRMSD and R_*ee*_. Table S1: Comparison of chemical properties of different metal ions. Table S2: Maximum ISO values of the isosurfaces obtained from spatial density map analysis.

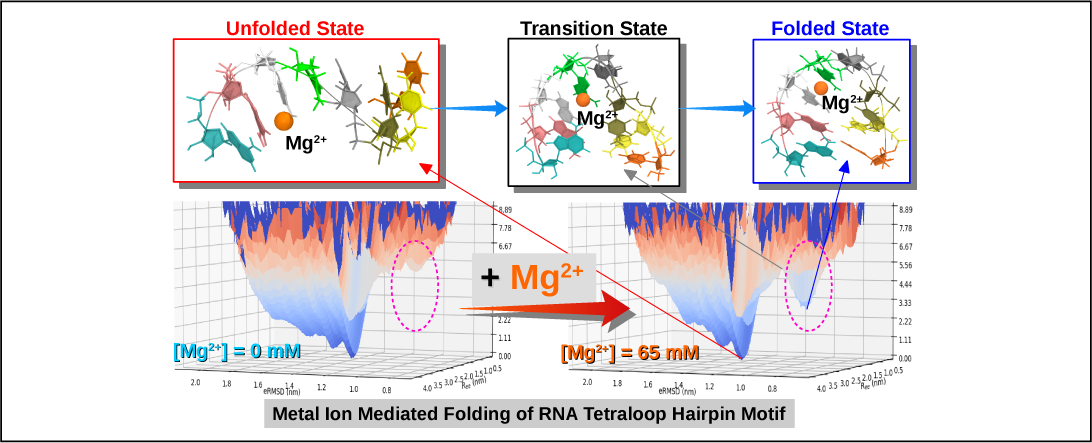
Graphical TOC Entry

