## Supplementary Information for "Mg^2+^ Sensing by an RNA Fragment: Role of Mg^2+^ Coordinated Water Molecules"

### Supporting Information for "$Mg^{2+}$ Sensing by an RNA Fragment: Role of $Mg^{2+}$ Coordinated Water Molecules"

**Calculation of coordination number between two sets of atoms** The COORDINATION keyword of PLUMED package[1,2] is used to calculate coordination number between two groups of atoms (say group A & group B) over a given MD trajectory. At each time step, we calculate the switching function $S_{ij}$ where,

$$S_{ij} = \frac{1 - (\frac{r_{ij}}{r_0})^6}{1 - (\frac{r_{ij}}{r_0})^{12}} \tag{1}$$

In the above function, $r_{ij}$ represents the inter-atomic distances between the atoms of group A (indexed with $i$) and atoms of group B (indexed with $j$), and $r_0$ represents the cut-off distance. Finally the coordination number (CN) is calculated as,

$$CN = \sum_{i \in A} \sum_{j \in B} S_{ij}. \tag{2}$$

We calculate the coordination number between metal ions with respect to some key nucleosides to estimate whether a metal ion is present inside the tetraloop or outside. Crystal structure[3] of the L4 hairpin motif suggests that formation of a metal ion mediated base pair between the terminal residues of the tetraloop (C69 and A72) is essential for its folding. This pair is followed by the C68:G73 base pair. So we add all the metal ions present in the system in group A and add the nucleobase and sugar atoms of the four key nucleotides, C68, C69, A72 and G73 in group B. Since, the $[Mg^{2+}]$ = 65 mM system contains both $K^+$ and $Mg^{2+}$ ions, we further calculate $CN_K$ and $CN_{Mg}$ separately where the group A contains only the $K^+$ ions (the former case) or only the $Mg^{2+}$ ions (the later case). Similarly, for the $[Ca^{2+}]$ = 65 mM system that contains both $K^+$ and $Ca^{2+}$ ions, we further calculate $CN_K$ and $CN_{Ca}$ separately where the group A contains only the $K^+$ ions (the former case) or only the $Ca^{2+}$ ions (the later case). The cut-off distance ($r_0$) is set to 5 nm.

**Parameters used for calculating the electrostatic potential surfaces in the PBEQ Solver server**

1. **Dielectric constant for the reference environment:** $\varepsilon_R = 1.0$

2. **Dielectric constant for the RNA interior:** $\varepsilon_P = 1.0$

3. **Solvent dielectric constant:** $\varepsilon_W = 80$

4. **Salt concentration:** Conc = 0.15 (mole/liter)

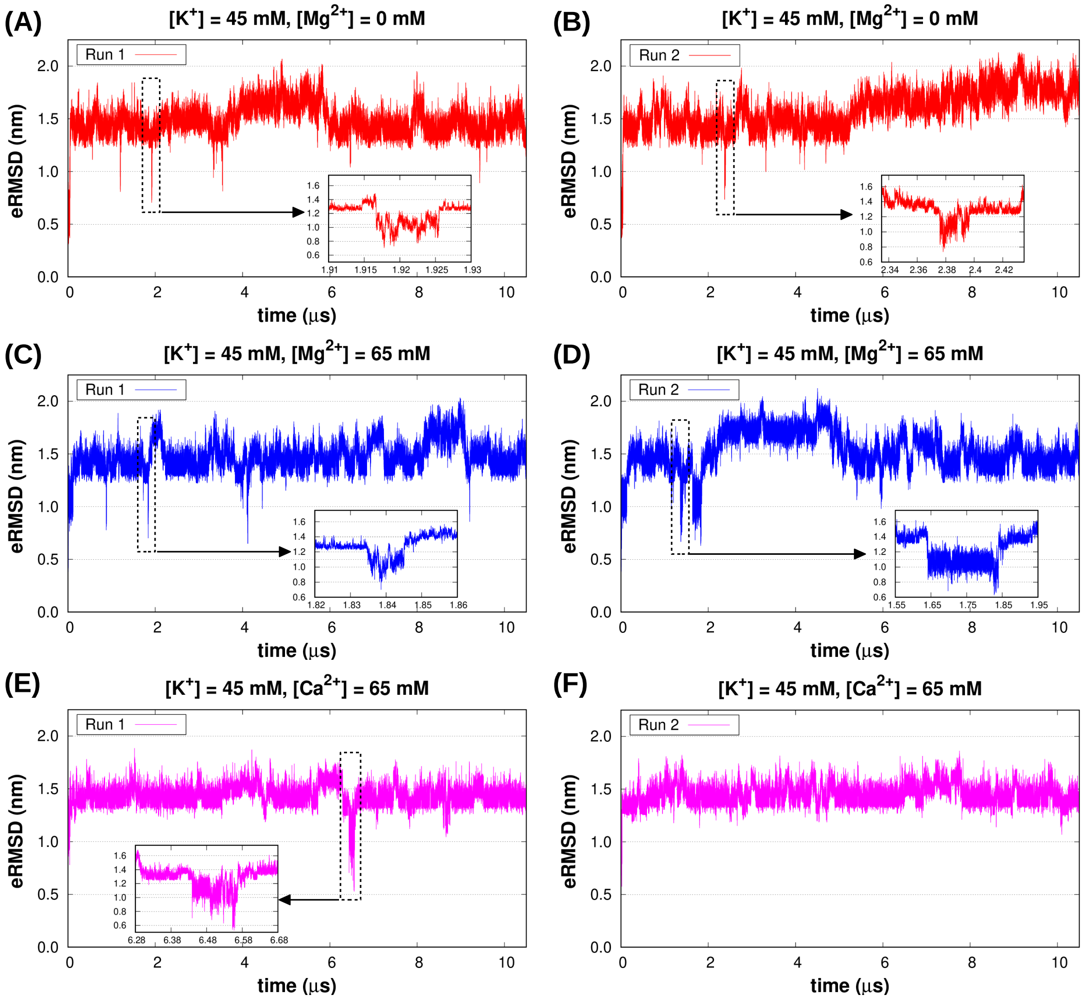


Figure S1: eRMSD values of the RNA fragment with respect to the energy minimized structure are calculated over the 10.5 $\mu$s long trajectories obtained at (A,B) $[K^+] = 45$ mM, $[Mg^{2+}] = 0$ mM; (C,D) $[K^+] = 45$ mM, $[Mg^{2+}] = 65$ mM and (E,F) $[K^+] = 45$ mM, $[Ca^{2+}] = 65$ mM. In each trajectory (except F) one unfolded$\rightarrow$folded$\rightarrow$unfolded transition is zoomed in and shown in the inset. For each trajectory the initial 0.5 $\mu$s data were ignored and the rest 10 $\mu$s were used for analysis.

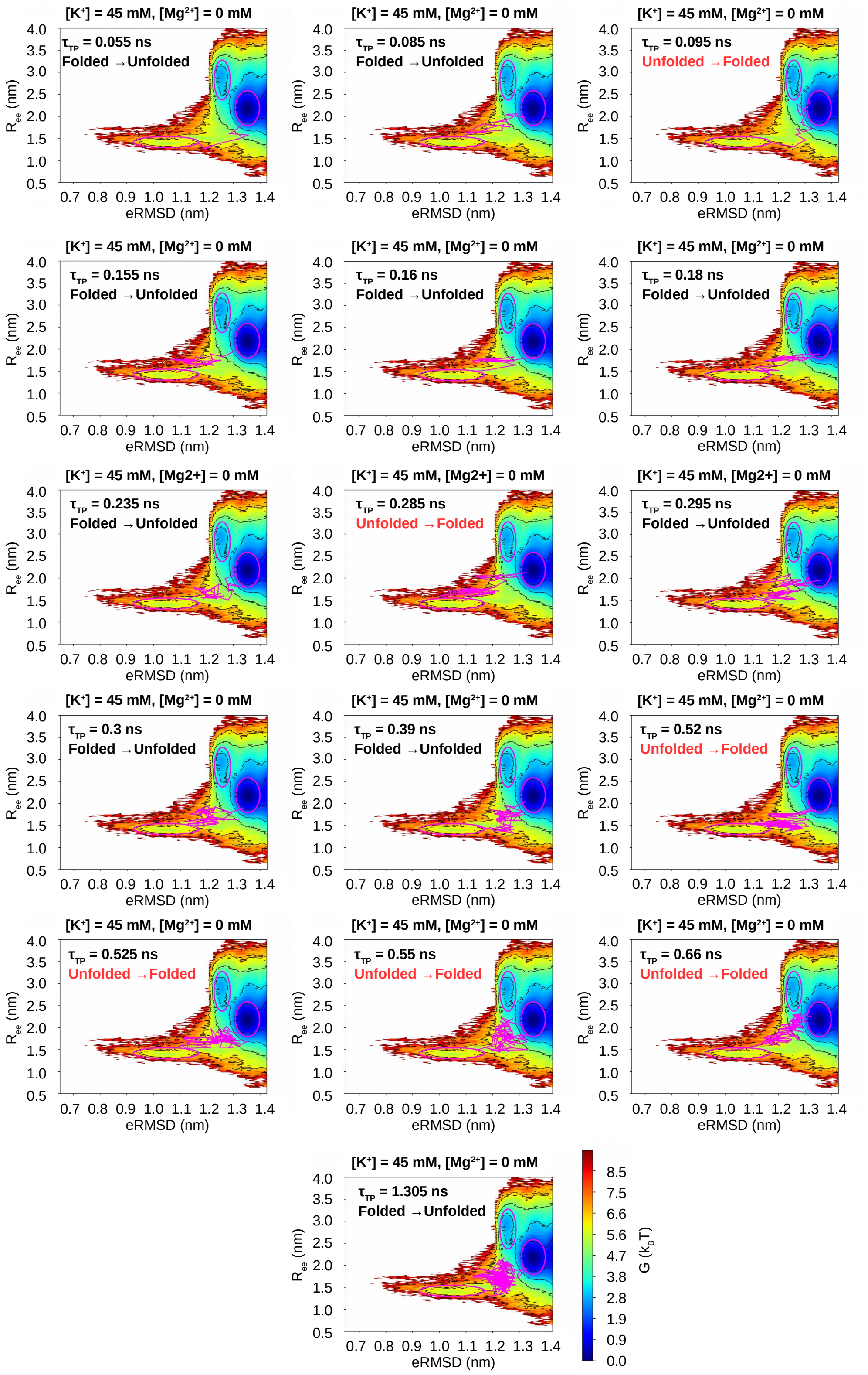


Figure S2: Transition paths (shown in magenta color) connecting the folded and unfolded basins for the system corresponding to $[Mg^{2+}] = 0$ mM. $\tau_{TP}$ represents the time length of each transition path (TP).

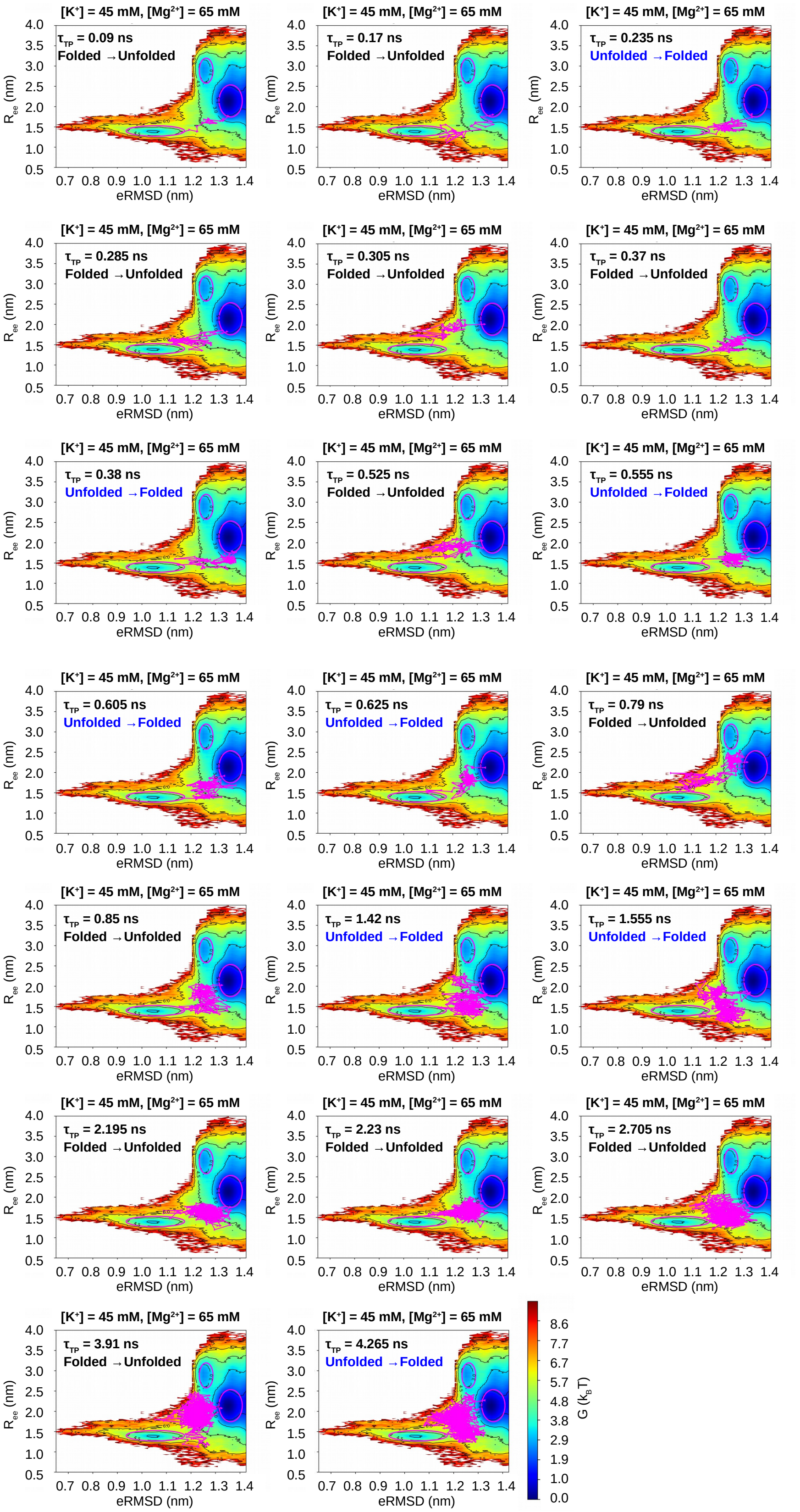


Figure S3: Transition paths (shown in magenta color) connecting the folded and unfolded basins for the system corresponding to [$Mg^{2+}$] = 65 mM. $\tau_{TP}$ represents the time length of each transition path (TP).

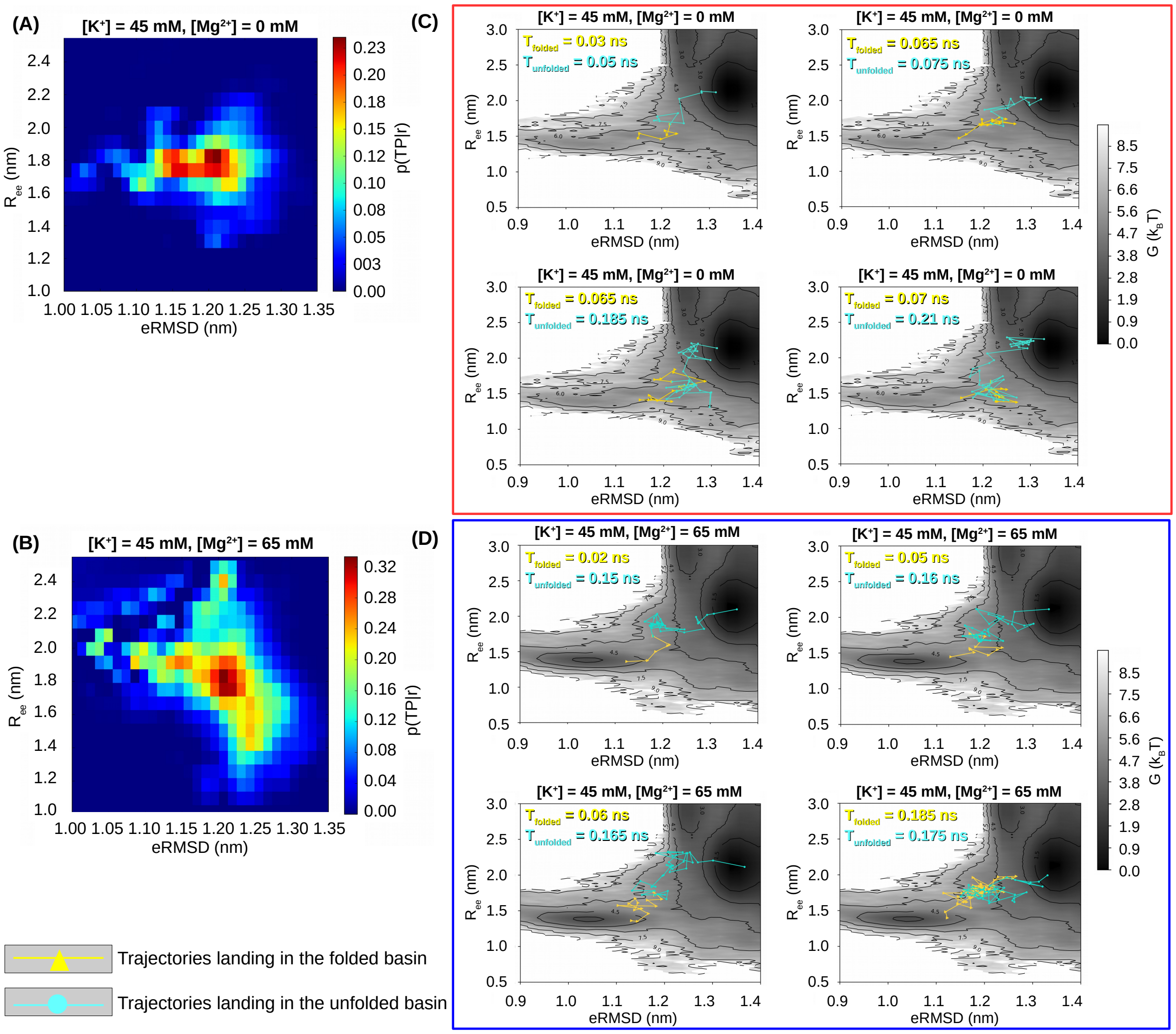


Figure S4: Distribution of the probability p(TP|$r$) in the configuration space (defined by eRMSD and $R_{ee}$) is shown for (A) $[Mg^{2+}]$ = 0 mM and (B) $[Mg^{2+}]$ = 65 mM. The maximum value of p(TP|$r$) is observed at eRMSD = 1.2 nm and $R_{ee}$ = 1.8 nm for both the cases. Out of all the $P_{fold}$ trajectories, 4 representative trajectories that land in the folded basin and 4 representative trajectories that land in the unfolded basin are shown for (C) $[Mg^{2+}]$ = 0 mM and (D) $[Mg^{2+}]$ = 65 mM.

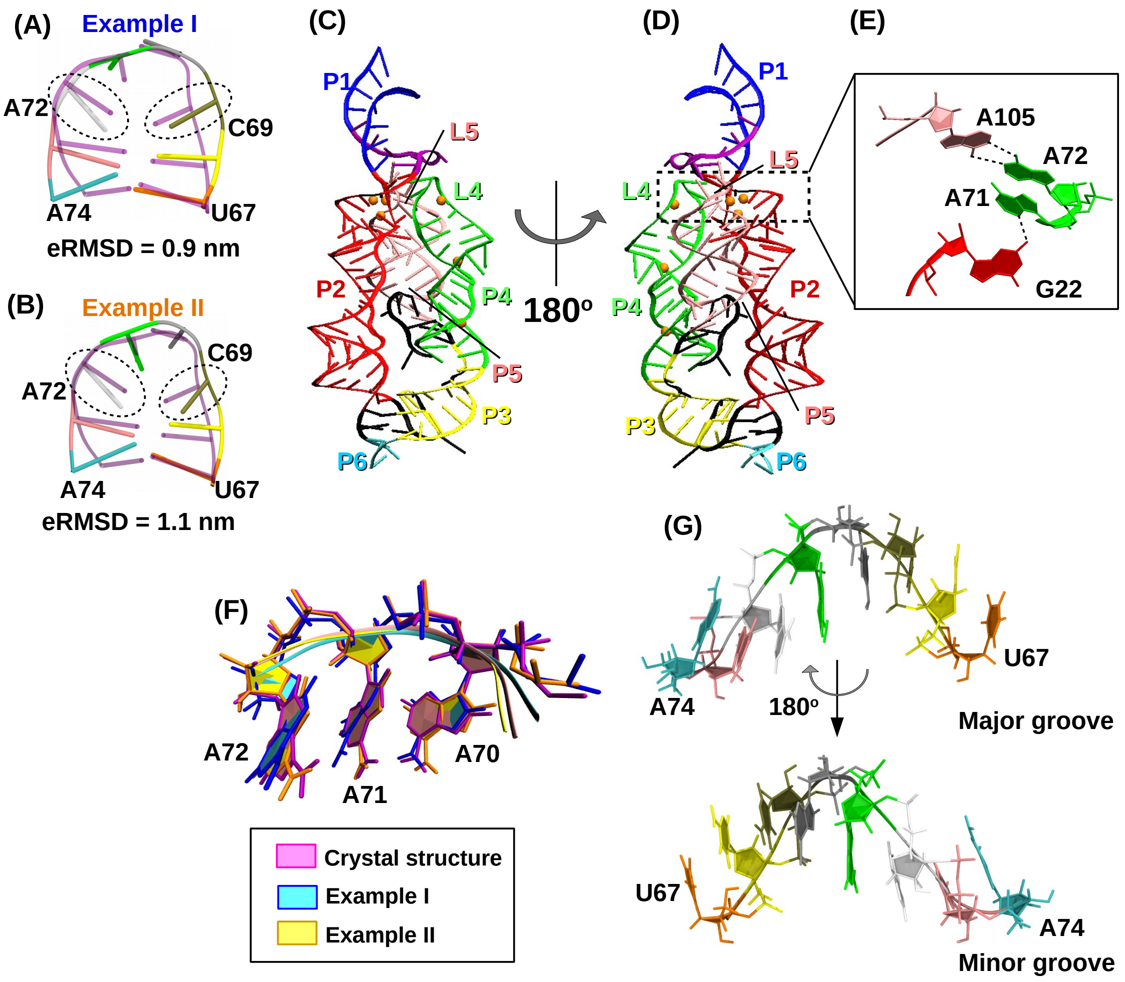


Figure S5: (A, B) Examples of RNA conformations that populate the folded basin centered at eRMSD $\approx$ 1.04 nm are superposed over the crystal structure. The crystal structure is shown in transparent cartoon representation in magenta. Regions of major changes are encircled. The corresponding eRMSD values are reported. One example has (A) eRMSD $<$ 1.04 nm and the other example has (B) eRMSD $>$ 1.04 nm. (C) Structure of the complete 'M-box' riboswitch (PDB Id: 2QBZ) is shown in cartoon representation. Different secondary structural motifs (P1, P2, P3, P4, P5, P6, L4 and L5) are represented in different colors. (D) Same molecule is shown after a 180° rotation. (E) Hydrogen bonding interactions between A71 with G22 (P2 stem) and A72 with A105 (L5 loop) are shown in broken lines. (F) Conservation of the consecutive stacking interactions between A70, A71 and A72 bases in the folded conformation are illustrated by superposing folded structures (Example I & Example II) on the crystal structure. (G) Illustration of the single stranded helix like unfolded structure from the major groove side and minor groove side. Nucleobases from all the eight residues are found to be sequentially stacked.

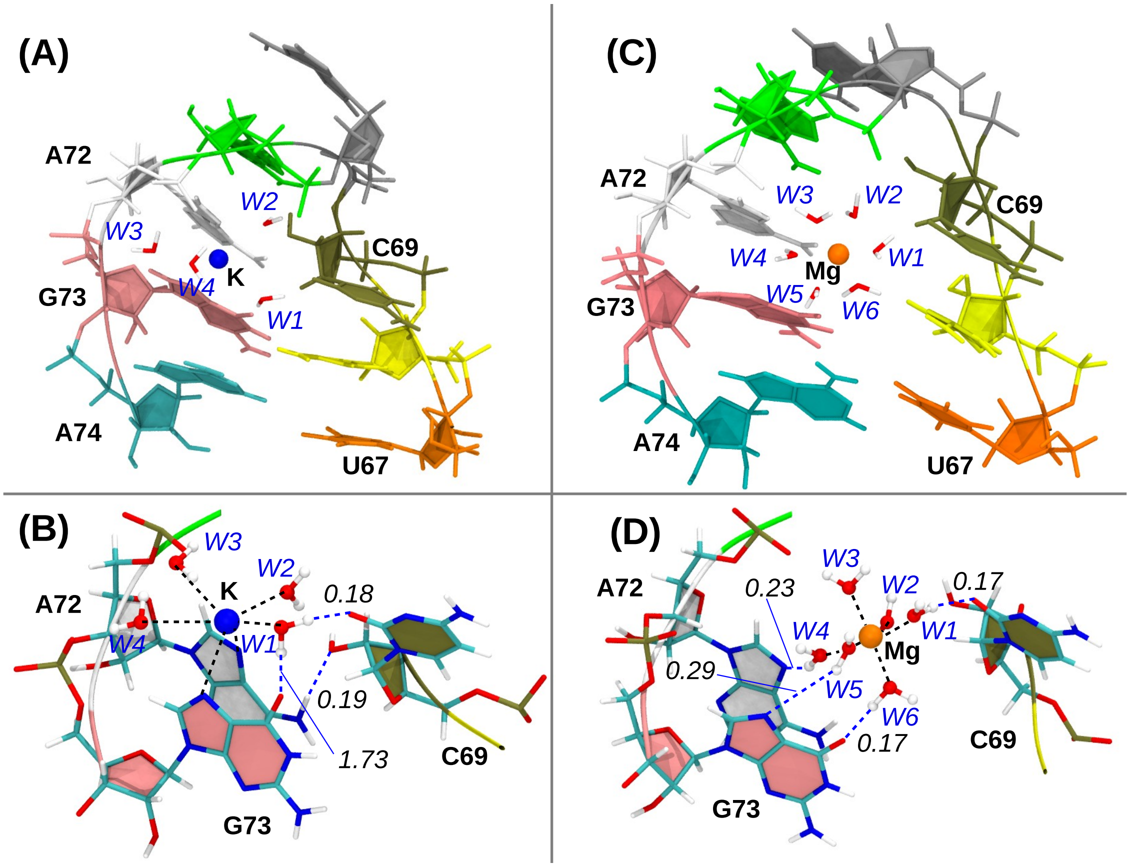


Figure S6: (A) Folded conformation of the L4 tetraloop hairpin motif when a $K^+$ is bound at its center. The 4 water molecules present in the first coordination shell of the ion are labeled as W1, W2, W3 and W4. (B) Details of the $K^+$-RNA interactions are highlighted. The $K^+$ directly interacts with four water molecules and N7 atoms of both A72 and G73. These contacts are shown in black broken line. Two hydrogens of the water W1 form two different hydrogen bonds with O6 of G73 and O2 of C69. There is also a hydrogen bond between N6 of A72 and O2′ of C69. These hydrogen bonds are shown as blue broken lines. (C) Folded conformation of the hairpin when a $Mg^{2+}$ is bound at its center. The 6 water molecules present in the first coordination shell of $Mg^{2+}$ are labeled as W1, W2, W3, W4, W5 and W6. (D) Details of the $Mg^{2+}$-RNA interactions are illustrated. Interaction between $Mg^{2+}$ and 6 water molecules are shown in black broken lines. The waters W4, W5 and W6 form hydrogen bonds with N7 of A72, N7 of G73 and O6 of G73, respectively. Another water W1 forms a hydrogen bond with O2 of C69. All these hydrogen bonds are shown in blue broken lines. For each hydrogen bond, the corresponding distance between the hydrogen and the acceptor is reported in nm. The ions are represented as spheres with radii equal to 30% of their respective van der Waals radius.

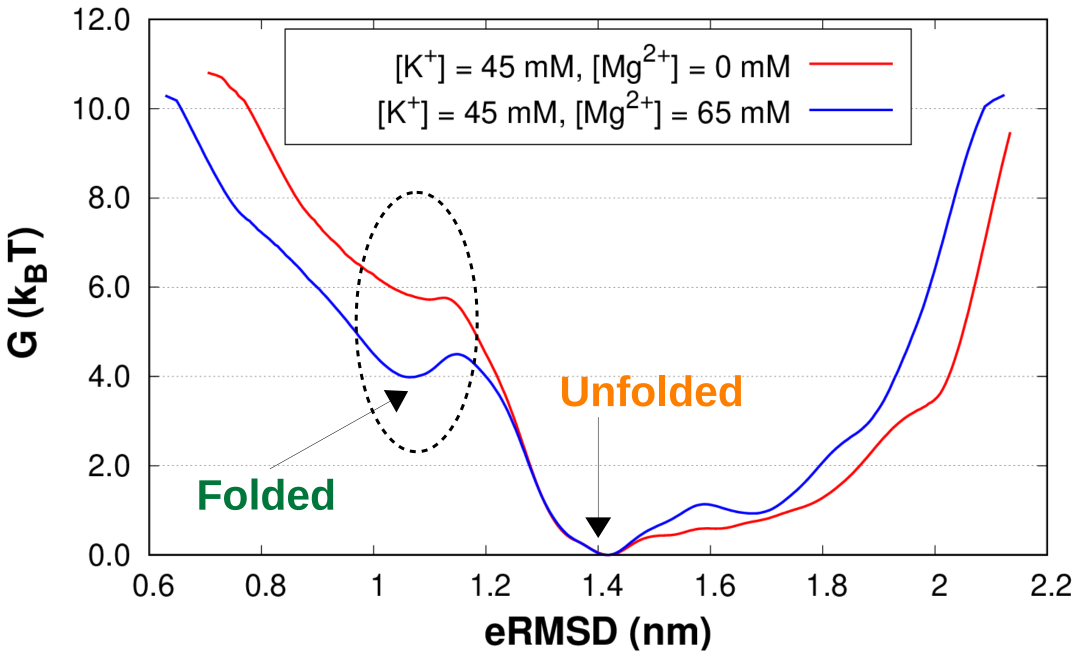


Figure S7: FES of L4 tetraloop folding projected onto eRMSD at two different ion concentrations – (red) $[K^+] = 45$ mM, $[Mg^{2+}] = 0$ mM and (blue) $[K^+] = 45$ mM, $[Mg^{2+}] = 65$ mM. $Mg^{2+}$ ions stabilize the folded state.

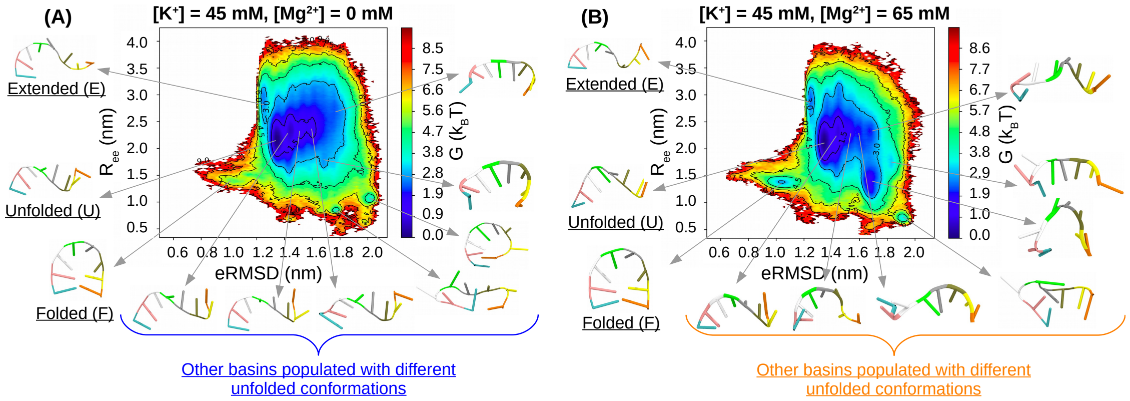


Figure S8: FES of L4-tetraloop folding projected onto the CVs, eRMSD and $R_{ee}$ for the systems (A) $[Mg^{2+}] = 0$ mM and (B) $[Mg^{2+}] = 65$ mM. The FES shows 3 major basins: (1) Folded (eRMSD $\approx$ 1.04 nm, $R_{ee} \approx$ 1.38 nm), (2) Extended (eRMSD $\approx$ 1.25 nm, $R_{ee} \approx$ 2.90 nm) and (3) Unfolded (eRMSD $\approx$ 1.35 nm, $R_{ee} \approx$ 2.15 nm). In addition to these there are seven other basins populated with different unfolded conformations of RNA (eRMSD $\geq$ 1.4 nm). Representative structures for each basin are shown in cartoon representation.

Table S1: Comparison of the chemical properties of $Mg^{2+}$ with common biological ions. The table is adopted from work by Maguire and Cowan.[4]

| Ion | Ionic radius[5,6] (Å) | Hydrated radius[5] (Å) | Ratio of radii | Ionic volume ($Å^3$) | Hydrated volume ($Å^3$) | Ratio of volumes | Coordination number | Water exchange rate[5] ($sec^{-1}$) | Transport number[7] |
|---|---|---|---|---|---|---|---|---|---|
| $Na^+$ | 0.95 | 2.75 | 2.9 | 3.6 | 88.3 | 24.5 | 6 | $8 \times 10^8$ | 7 – 13 |
| $K^+$ | 1.38 | 2.32 | 1.7 | 11.0 | 52.5 | 4.8 | 6–8 | $10^9$ | 4 – 6 |
| $Ca^{2+}$ | 0.99 | 2.95 | 3.0 | 4.1 | 108 | 26.3 | 6–8 | $3 \times 10^8$ | 8 – 12 |
| $Mg^{2+}$ | 0.65 | 4.76 | 7.3 | 1.2 | 453 | 394 | 6 | $10^5$ | 12 – 14 |

Table S2: Maximum ISO values of the metal ions obtained from spatial density map analysis illustrated in Figure 5.

| | $[Mg^{2+}] = 0$ mM | $[Mg^{2+}] = 65$ mM | | |
|---|---|---|---|---|
| Basin | $K^+$ | only $K^+$ | Only $Mg^{2+}$ | Both $K^+$ and $Mg^{2+}$ |
| Folded | 0.65 | 0.18 | 0.50 | 0.60 |
| Unfolded | 0.44 | 0.19 | 0.18 | 0.19 |
| Extended | 0.60 | 0.06 | 0.25 | 0.26 |
